# Soybean Root Nodule Occupancy: Competition Between *Bradyrhizobium* and *Sinorhizobium* Strains Inoculated at Different Plant Growth Stages

**DOI:** 10.1101/2025.10.27.684879

**Authors:** Matthew Knoll, Babur S Mirza

## Abstract

Soybean is frequently nodulated by species from the *Bradyrhizobium* (BR) and/or *Sinorhizobium* (SR) genera. Several factors, such as soil pH, host genotype, geographic location, and other environmental variables, are reported to influence the preferential selection between BR or SR species within soybean root nodules. However, it remains unclear whether the age of the host plant at the time of inoculation affects preferential rhizobial selection. To investigate this, we inoculated soybean plants with different cell densities of BR and SR strains at three time points: at sowing (T_0_), two weeks after germination (T_2_), and four weeks after germination (T_4_). We used 16S rRNA gene amplicon sequencing of root nodule and rhizosphere samples to assess the relative abundance of BR and SR in nodule and rhizosphere. We observed a clear shift in nodule occupancy that favored BR at the time of seed sowing (T_0_) but increasingly favored SR when plants were inoculated at T_2_ and T_4_ stages. Specifically, at T_4_, SR dominated in nodules across all treatments, representing 88–99% of total sequences, regardless of applied inoculum ratio. In contrast, a similar number of sequences for both strains were detected in the rhizosphere at the time of the final harvest. These results highlight host age as an important ecological driver in legume–rhizobium interactions and suggest that inoculation time strongly influences microsymbiont selection. This information is important in understanding rhizobial competition and optimizing the timing of inoculation for soybean.

**IMPORTANCE:** Soybean is one of the world’s most valuable crops and fulfills most of its nitrogen requirements by developing symbiotic associated with nitrogen-fixing rhizobia. This reduces the need for chemical fertilizers by converting atmospheric nitrogen into a plant useable form of nitrogen. Multiple species from four rhizobial genera can nodulate soybean, and the plant’s choice of rhizobial partner is reported to change depending on environmental conditions such as pH, host genotype, geographic location, and other environmental factors. This study explores how the age of the soybean plant affects its preference for two frequently reported beneficial rhizobial species (*B. diazoefficiens* and *S. fredii*). By testing inoculation at different growth stages, we discovered that at early growth stages plants favored BR, while older plants increasingly selected SR for nodule formation. These findings highlight the level of complexity in plant–microbe interactions and could help optimize bioinoculant strategies for improving sustainability and crop yields.

## INTRODUCTION

Soybean (*Glycine max* L.) is a globally important legume that fulfills most of its nitrogen requirements through symbiotic associations with nitrogen-fixing rhizobia (1). Four rhizobial genera including; *Bradyrhizobium, Sinorhizobium* (formerly *Ensifer*), *Rhizobium*, and *Mesorhizobium* are known to nodulate soybean (2-14). Among these, *Bradyrhizobium* and *Sinorhizobium* are the most agriculturally significant and are frequently reported to occupy soybean root nodules worldwide. Although these two genera are genetically distinct and markedly differ in physiological attributes, both are capable of efficiently nodulating soybean plants. For instance, *Bradyrhizobium* species typically possess larger genomes (e.g., ∼9.1 Mb) and are slow-growing, with doubling times ranging from 7 to 10 hours (15). In contrast, *Sinorhizobium* species generally have smaller genomes (∼6.7 Mb) and exhibit faster growth with doubling times of only 1 to 2 hours (15). Despite these differences, species from both genera are equally effective in nodulating soybean and have been the focus of numerous competition-based studies (10, 11, 16-21).

Overall, preferential selection of specific rhizobial species among multiple host-compatible species in the rhizosphere is a complex process driven by rhizobial diversity, host genotype, and environmental factors (10, 17, 18, 22). In soybean, several studies have suggested that competition for nodulation between *Bradyrhizobium* and *Sinorhizobium* species is influenced by factors such as soil pH (4, 11, 17, 18, 23-25), nutrient availability and salinity (6, 26), land use (10), temperature (22, 27), host genotype (17, 24, 27) and diversity of non-rhizobial bacteria (16). In particular, *Bradyrhizobium* species are reported to dominate in acidic soils (5, 18, 24), whereas *Sinorhizobium* species are favored under alkaline conditions (4, 6-8, 12, 16).

Although the dynamics of nodule occupancy between *Bradyrhizobium* and *Sinorhizobium* species have been well studied, most studies have focused on early inoculation at the time of seed sowing or relied on natural soil rhizobial populations (10, 11, 16-21, 26, 28, 29). Root exudate composition in plants changes significantly during the first few weeks, shifting from simple sugars in early stages to an increased secretion of secondary compounds, such as flavonoids, isoflavonoids, phenolics, and amino acids, as plants mature (30-32). These previously demonstrated age-related changes in root exudates and signaling molecules are suggested to alter overall microbial community structure in the rhizosphere (16, 32, 33) and may also affect the competitive dynamics of nodule occupancy between *Bradyrhizobium* and *Sinorhizobium*. However, limited information is available on the relative competitiveness of *Bradyrhizobium* and *Sinorhizobium* for nodulation in relation to the age of host.

Hence, the primary objective of this study was to investigate the effect of soybean plant age on the competition between two well-characterized rhizobial strains—*Bradyrhizobium diazoefficiens* USDA 110 (BR) and *Sinorhizobium fredii* USDA 191 (SR). We conducted a greenhouse study in which soybean plants were inoculated with two different strains either individually or co-inoculated at three growth stages: at seed sowing, and at two and four weeks after planting. For the co-inoculation treatments, we applied varying inoculum ratios of BR:SR (1:1, 1:100, and 100:1) at each growth stage. To assess strain-specific nodule occupancy and rhizosphere colonization, we analyzed the microbiomes of root nodules and rhizosphere microbes from these plants using Illumina paired-end DNA sequencing of 16S rRNA amplicons. At harvesting (8 weeks of plant growth), we also measured plant physiological parameters such as nodule number, and shoot and root biomass, to evaluate the potential effects of rhizobial strain selection on soybean growth.

## RESULTS AND DISCUSSION

Overall, we retrieved 9.82 million high-quality 16S rRNA sequences across all samples, including 3.94 million sequences from root nodules of 71 co-inoculated soybean plants (10 nodules per plant), 3.74 million sequences from 24 control plants inoculated with either BR or SR strains at three different growth stages, and 2.13 million sequences from the rhizosphere samples. All sequences from root nodules were associated with *B. diazoefficiens* USDA 110 or *S. fredii* USDA 191, and were 100% identical to the inoculated strains. In contrast, only 3.2% of rhizosphere sequences (68,123 out of 2,135,068 total sequences) were identical to BR or SR strains. The rhizosphere sequences were predominantly related to *Bacillus* spp. (64%), along with a few other genera, each comprising approximately 4–7% of the total sequences. Lastly, we did not observe any nodulation in uninoculated control plants. In addition, insufficient DNA (<1 ng/µl) was recovered from the no-plant controls (sand without plants inoculated with rhizobia), and PCR amplification was not successful. A similar culture–independent approach of using 16S rRNA gene amplicon sequencing has been adopted to study rhizobial communities within soybean root nodules (14, 34-37).

### Distribution of 16S rRNA gene sequences in root nodules of control plants inoculated with either BR or SR strains

Control plants inoculated with either BR or SR at three different growth stages developed successful nodulation with their respective strains. Nearly all the sequences (3.74 million) retrieved from these nodules corresponded to the inoculated strain (**Fig. 1**). For example, in BR-inoculated plants, BR-related sequences dominated (>99.3%) across all three inoculation times (T_0_, T_2_, and T_4_). Similarly, SR-inoculated plants showed a high abundance (96.5–99.9%) of SR-related sequences at all time points and this was consistent for both overall sequences (**Fig. 1; Table 1)** and the normalized data adjusted for the three copies of 16S rRNA gene per SR cell **(Fig. S1; Table 1**). These results indicate that both rhizobial strains are compatible with soybean, consistent with previous studies (10, 11, 16-21, 26, 28, 29), and can effectively nodulate soybean at different stages of plant growth. Although the dominant sequences aligned with the inoculated strain, a small fraction (mostly <1%) of 16S rRNA gene sequences related to the non-inoculated strain was also detected. For instance, in plants inoculated with BR, SR-related sequences accounted for only 0.3% (T_0_), 0.26% (T_2_), and 0.23% (T_4_). Similarly, BR-related sequences were present at low abundance in root nodules of SR inoculated plants (**Fig. 1; Fig. S1**). These low-frequency contaminated sequences may result from index hopping or barcode misassignment, as previously reported in ∼2% of reads from mixed-sample analyses using Illumina platform (38). Alternatively, minor cross-contamination via aerial transmission from neighboring plants or greenhouse soil is also a possibility. Nevertheless, these results confirm that each strain maintained high purity in nodule colonization when inoculated individually and that soybean plants successfully formed nodules with either strain across all inoculation time points.

**Table 1.** Parameters of soybean plants inoculated with *Bradyrhizobium diazoefficiens* USDA 110 (BR) and *Sinorhizobium fredii* USDA 191 (SR), either alone or in combination at different cell ratios. Reported parameters include BR:SR sequences ratio, number of plants, total number of sequences, percentage of BR and SR sequences, percentage of BR and SR sequences (adjusted copies), average BR sequences, average SR sequences, average number of nodules per plant, shoot weight, and root weight. Values are reported as means with standard error (±SE).

| Inoculation Time | Treatment | BR:SR Ratio | Plants (n) | Total Seqs | BR:SR Seqs (%) | Adj. BR:SR Seqs (%) | BR Seqs (Avg.) | SR Seqs (Avg.) | Nodules/plant (Avg.) | Shoot wt | Root wt |
| --- | --- | --- | --- | --- | --- | --- | --- | --- | --- | --- | --- |
| T0 | control | 1:0 | 2 | 131,240 | 99:1 | 100:0 | 19,341±13,717 | 641±3 | 25±3 | 4.19±0.07 | 2.35±0.06 |
| | control | 0:1 | 2 | 125,009 | 3:97 | 9:91 | 2,163±2,020 | 60,341±8,378 | 20±3 | 3.4±0.14 | 2.24±0.01 |
| | Co-inocula | 1:1 | 7 | 203,996 | 89:11 | 96:4 | 25,956 ± 2,983 | 3,186 ± 2,236 | 31±2 | 4.18±0.11 | 2.04±0.05 |
| | Co-inocula | 1:100 | 7 | 212,507 | 15:85 | 35:65 | 4,674 ± 1904 | 25,684 ± 5,433 | 25±2 | 3.83±0.22 | 1.96±0.03 |
| | Co-inocula | 100:1 | 8 | 157,853 | 90:10 | 97:3 | 17,832 ± 3,311 | 1,900 ± 583 | 19±2 | 3.84±0.12 | 2.12±0.11 |
| T2 | control | 1:0 | 5 | 392967 | 99:1 | 99:1 | 129,970±49,040 | 1,019±335 | 36±5 | 3.85±0.09 | 2.27±0.02 |
| | control | 0:1 | 3 | 1047552 | 0:100 | 0:100 | 276±43 | 209,235±36,044 | 25±2 | 3.23±0.15 | 2.21±0.02 |
| | Co-inocula | 1:1 | 9 | 300,945 | 7:93 | 19:81 | 2,369 ± 1,431 | 31,070 ± 7,243 | 30±3 | 3.56±0.17 | 1.95±0.03 |
| | Co-inocula | 1:100 | 6 | 774,632 | 4:96 | 11:89 | 5,278 ± 5,125 | 123,827 ± 23,797 | 23±3 | 2.74±0.01 | 2.13±0.07 |
| | Co-inocula | 100:1 | 9 | 565,571 | 51:49 | 76:24 | 31,925 ± 16,568 | 30,916 ± 13,891 | 25±5 | 3.0±0.17 | 2.05±0.06 |
| T4 | control | 1:0 | 6 | 877,186 | 99:1 | 99:1 | 145,200±18,127 | 998±339 | 27±3 | 3.45±0.07 | 2.34±0.03 |
| | control | 0:1 | 6 | 1,166,064 | 0:100 | 0:100 | 528±274 | 193,816±50,966 | 23±5 | 3.25±0.19 | 2.38±0.03 |
| | Co-inocula | 1:1 | 9 | 745,141 | 3:97 | 7:93 | 2,132 ± 1,526 | 80,662 ± 1,4604 | 20±3 | 3.03±0.11 | 2.17±0.13 |
| | Co-inocula | 1:100 | 7 | 387172 | 12:88 | 29:71 | 6,557 ± 3,062 | 48,754 ± 13,229 | 19±6 | 3.35±0.25 | 2.14±0.04 |
| | Co-inocula | 100:1 | 9 | 593,408 | 7:93 | 19:81 | 4,723 ± 2,753 | 61,211 ± 9,119 | 16±3 | 3.310.14 | 2.16±0.03 |

**Fig. 1.**
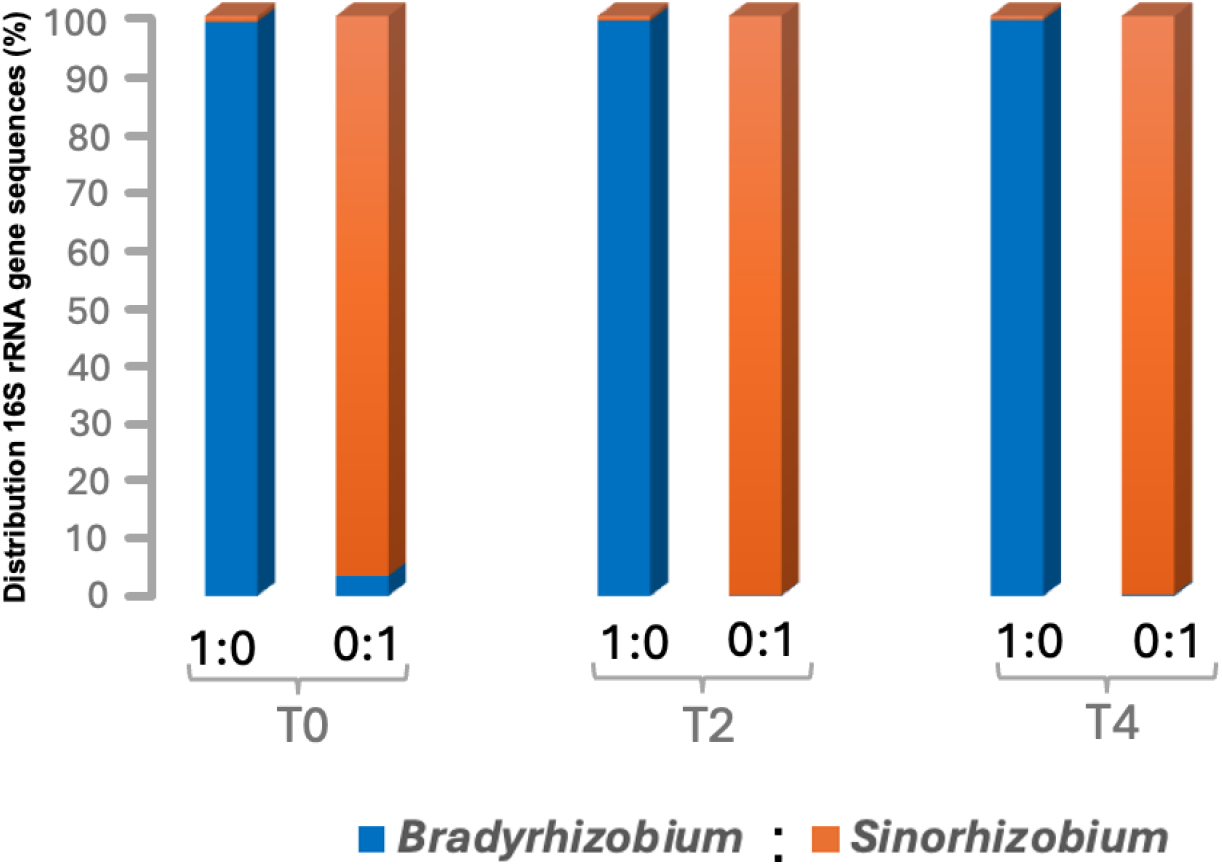
Relative distribution of *Bradyrhizobium diazoefficiens* USDA 110 (BR-blue) and *Sinorhizobium fredii* USDA 191 (SR - tan) within soybean root nodules from 24 control plants inoculated with either strain at three time points: sowing (T_0_), two weeks (T_2_), and four weeks (T_4_) after planting. A total of 3.74 million sequences were analyzed from 24 plants, with 10 nodules randomly selected per plant.

### Distribution of 16S rRNA gene sequences in root nodules of co-inoculated plants with BR and SR at three different stages of plant growth (T_0_, T_2_, and T_4_)

Most previous studies on competition between *Bradyrhizobium* and *Sinorhizobium* species for soybean nodulation have emphasized soil pH and host genotype as the primary determinants of their preferential selection. In acidic to neutral soils, *Bradyrhizobium* species typically dominate, whereas alkaline soils favor *Sinorhizobium* species (4, 11, 17-20, 23-25). Similarly, Asiatic soybean cultivars often show stronger compatibility with *Sinorhizobium* species, while American cultivars preferentially select *Bradyrhizobium* species (17, 24, 27). All of the above studies either inoculated soybean at the time of seed sowing or relied on natural rhizobial populations that were already present in soil before seed sowing. So far, little information is available on the potential influence of host plant age on rhizobial species selection.

We investigated whether the time of inoculation affects the preferential selection of rhizobial strains for soybean nodule occupancy. Soybean plants were inoculated with BR and SR strains at three time points: at sowing (T_0_), two weeks after germination (T_2_), and four weeks after germination (T_4_), using three different BR:SR cell ratios (1:1, 1:100, and 100:1). We observed a significant shift in nodule occupancy that favored BR in plants inoculated at the time of seed sowing (T_0_) but increasingly favored SR when plants were inoculated at T_2_ and T_4_ stages (**Fig. 2; Fig. S2; Table 1**). For instance, at T_0_ with an equal cell density (1:1 ratio), BR-related sequences accounted for 89% (25,956 ± 2,983; average sequences per plant ± SE) of total nodule sequences, compared to 11% for SR (3,186 ± 2,236). At unequal cell densities, the nodule occupancy was mostly dominated by the strain present at higher density. As an example, at 1:100 BR accounted for 15% sequences (4,674 ± 1,904) versus SR with 85% (25,684 ± 5,433), whereas at 100:1 BR accounted for 90% (17,832 ± 3,311) of sequences, while SR sequences were only 10% (1,900 ± 583) (**Fig. 2; Fig. S2; Table 1**). These results indicate a clear dominance of the strain inoculated at higher concentration at T_0_ and BR dominated at equal cell density. Furthermore, we also observed a similar overall pattern of nodule occupancy for normalized data adjusted for 16S rRNA gene copy number for SR (three copies per genome) (**Supplemental Material and Fig. S3&S4**).

**Fig. 2.**
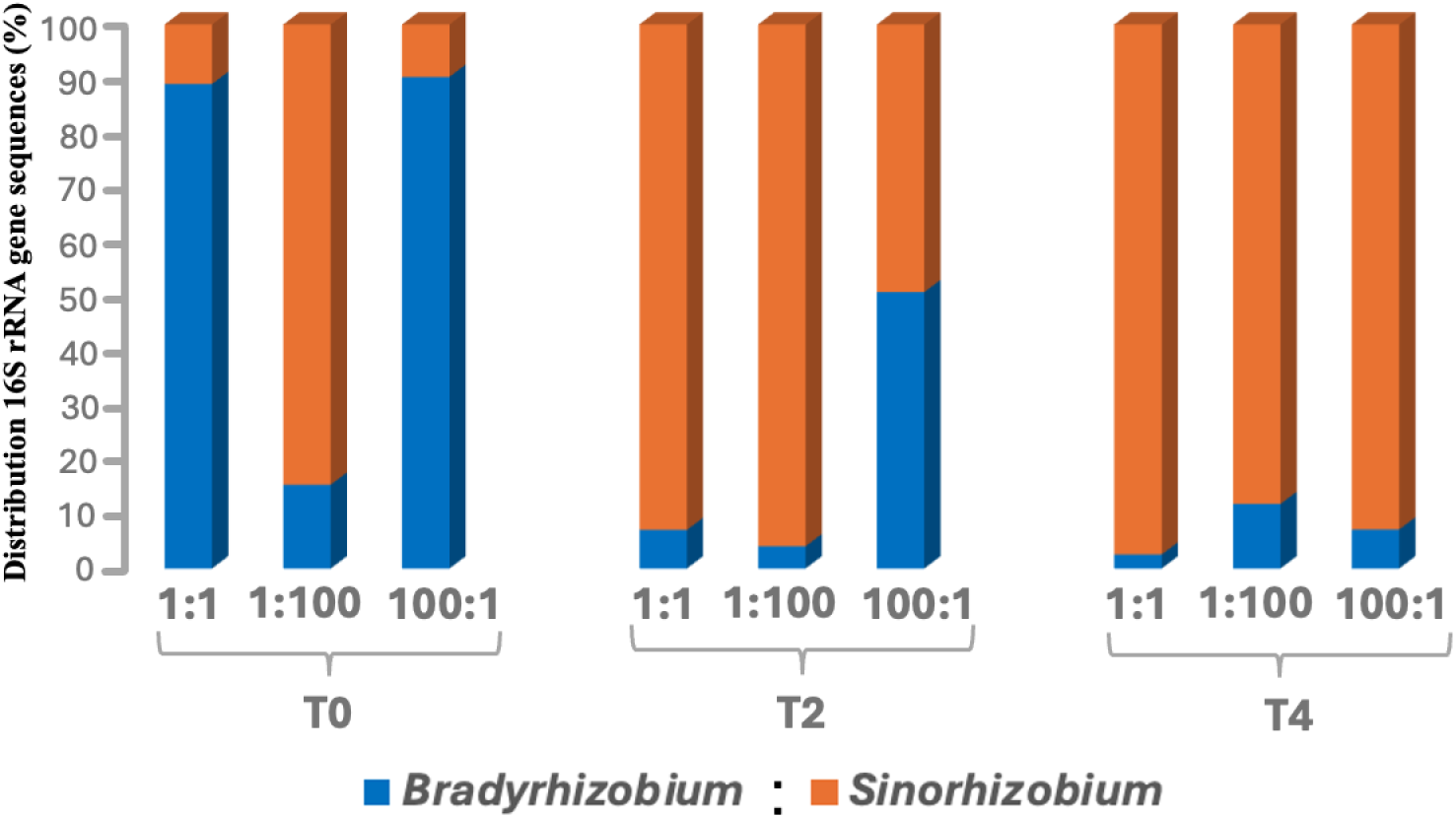
Relative distribution of *Bradyrhizobium diazoefficiens* USDA 110 (BR) and *Sinorhizobium fredii* USDA 191 (SR) within soybean root nodules of 71 plants co-inoculated with BR and SR at three time points: sowing (T_0_), two weeks (T_2_), and four weeks (T_4_) after germination, using inoculation three BR:SR ratios (1:1, 1:100, and 100:1). A total of 3.94 million sequences were analyzed from nodules (10 per plant).

Consistent with our results, previous studies have also demonstrated the preferential selection of *Bradyrhizobium* species over *Sinorhizobium* species under neutral pH conditions when soybean plants were co-inoculated with both strains at the time of seed sowing (11, 19). This early preferential selection of BR may be due to stronger initial adsorption of BR cells on soybean roots and/or better recognition of BR Nod factors during early plant growth stages (39, 40). Moreover, BR Nod factors have been reported to promote soybean seed germination and seedling growth at early stages, which may further bias partner choice toward BR (41).

Interestingly, the preferential selection of BR at the time of seed sowing was reversed when the plants were inoculated at later growth stage (**Fig. 2; Table 1**). In plants inoculated at T_2_, SR outcompeted BR for nodule occupancy. At a 1:1 ratio, SR comprised 93% (31,070 ± 7,243) of nodules compared to only 7% (2,369 ± 1,431) sequences for BR. Even under the 100:1 BR:SR inoculation, SR sequences were 49% (30,916 ± 13,891) relative to 51% (31,925 ± 16,568) for BR (**Fig. 2; Table 1**). This highlights the increased competitiveness and improved compatibility of SR with the host plant at later stages that can reduce the initial advantage of high cell density of BR in inoculum. When SR was applied at 100-fold higher density (1:100), SR overwhelmingly dominated with 96% (123,827 ± 23,797) compared to only 4% (5,278 ± 5,125) for BR (**Fig. 2; Table 1**). The preferential selection of SR at the T_4_ stage was even more pronounced, where SR became the primary nodule occupant across all inoculum ratios. At equal densities (1:1), SR represented 97% (80,662 ± 14,604), while BR declined to only 3% (2,132 ± 1,526). Even at the 100:1 BR:SR ratio, SR maintained dominance at 93% (61,211 ± 9,119) compared to 7% (4,723 ± 2,753) for BR (**Fig. 2; Table 1**). A similar trend was observed in the normalized dataset, where SR was corrected for its three 16S rRNA gene copies. BR dominated nodules at early inoculation T_0_, whereas SR became more competitive at later stages (T_2_ and T_4_) (Supplemental Material and **Fig. S3 & S4**).

The competition for nodulation by different strains altered when plants were inoculated at different growth stages emphasizes a critical role of host plant age in shaping the outcome of rhizobial competition. During early development, soybean plants preferentially supported BR for nodulation, consistent with previous reports of BR dominance in root nodules under neutral pH conditions (11, 19). However, as plants matured, SR became increasingly competitive, eventually dominating even when BR was applied at higher cell densities (**Fig. 2; Fig. S2-4**). Several factors may contribute to this age-dependent shift. First, root exudate composition changes significantly with plant age, transitioning from simple sugars in seedlings to higher concentrations of phenolics, amino acids, and secondary metabolites in older plants (30-33, 42). In soybean, Li et al. (32) demonstrated that uninoculated plants produced 36% more flavonoid-related compounds and 71% more phenolic acids at four weeks compared to two-weeks time points, corresponding to our T_2_ and T_4_ inoculation treatments. Because flavonoids act as key signaling molecules in rhizobia–host interactions (39, 43), these compositional shifts may favor SR recognition and nodulation at later growth stages. Second, phenolic acids can serve as carbon sources for rhizobia (44), and variation in their composition may selectively promote the growth of specific strains (45, 46). Third, younger soybean plants may exert stronger partner selection, preferentially selecting BR to ensure efficient nodulation under neutral pH conditions, whereas older plants with increasing nitrogen demands may relax this selective pressure, thereby enabling SR to establish nodules more effectively. Lastly, bacterial physiology may also contribute to this shift: BR appears better adapted to early soil–root environments, which facilitates nodulation in young seedlings, whereas SR benefits at later stages by exploiting the more complex exudate profile with its broader substrate range, faster growth, and greater metabolic flexibility (47).

It is important to note that we collected 10 root nodules per plant. Based on the current results, we are unable to determine whether the observed low abundance of BR or SR strains across different treatments is due to their presence in very few nodules, or whether both strains coexist within individual nodules at varying cell densities. In a previous study (14), we assessed the microbiome of 193 individual root nodules from nine field-grown soybean plants and observed that *Bradyrhizobium* was consistently the dominant rhizobial endophyte in each nodule (>95% of sequences), regardless of the size or location of the nodule on soybean roots.

### Distribution of rhizobial and other bacterial sequences in the rhizosphere of soybean plants co-inoculated with BR and SR strains

In contrast to root nodules, only a small number of sequences of the inoculated strains (68,173 in total) were retrieved from the rhizosphere, including 29,351 BR and 38,822 SR sequences. On average, BR and SR sequences in the rhizosphere ranged from 245 to 920 per sample at harvest, which was significantly lower compared to root nodules sequences (**Fig. 3; Table S1**). Overall, despite clear differences in nodule occupancy, the relative abundances of BR and SR sequences retrieved from the rhizosphere at final harvest remained consistent across different treatments. In the 1:1 treatment at T_0_, BR sequences were less abundant than SR in the rhizosphere (281 vs. 877); however, BR remained the dominant nodule endophyte (89% of root nodule sequences). Similarly, at T_2_ and T_4_, BR sequences increased relative to SR in some rhizosphere samples (e.g., T_2_ 1:1, BR 840 vs. SR 629; T_4_ 100:1, BR 380 vs. SR 334), whereas SR abundance increased in nodules. This decoupling between rhizosphere abundance and nodule occupancy suggests that nodule formation is not limited by strain availability (16) but is instead primarily driven by the host plant (14). This was more evident in the normalized data (adjusted for three copies of 16S rRNA per SR genome) for BR and SR cells in the rhizosphere (**Fig. S5**). It is important to note that the relative abundance and survival of BR and SR may differ between the time of inoculation or during plant growth and at the time of harvest. The quality and quantity of root exudates decrease significantly with plant age (48) which alters the relative abundances of rhizosphere bacterial species (48). Because rhizosphere sequences were analyzed only at final harvest stage (Day 56), temporal shifts in rhizobial strains during earlier growth stages cannot be excluded. Furthermore, our results indicate that sequences related to both strains were retrieved from the rhizosphere for all treatments, suggesting that shifts in nodule occupancy are more likely driven by the host plant rather than rhizobial relative abundance in the rhizosphere.

**Fig. 3.**
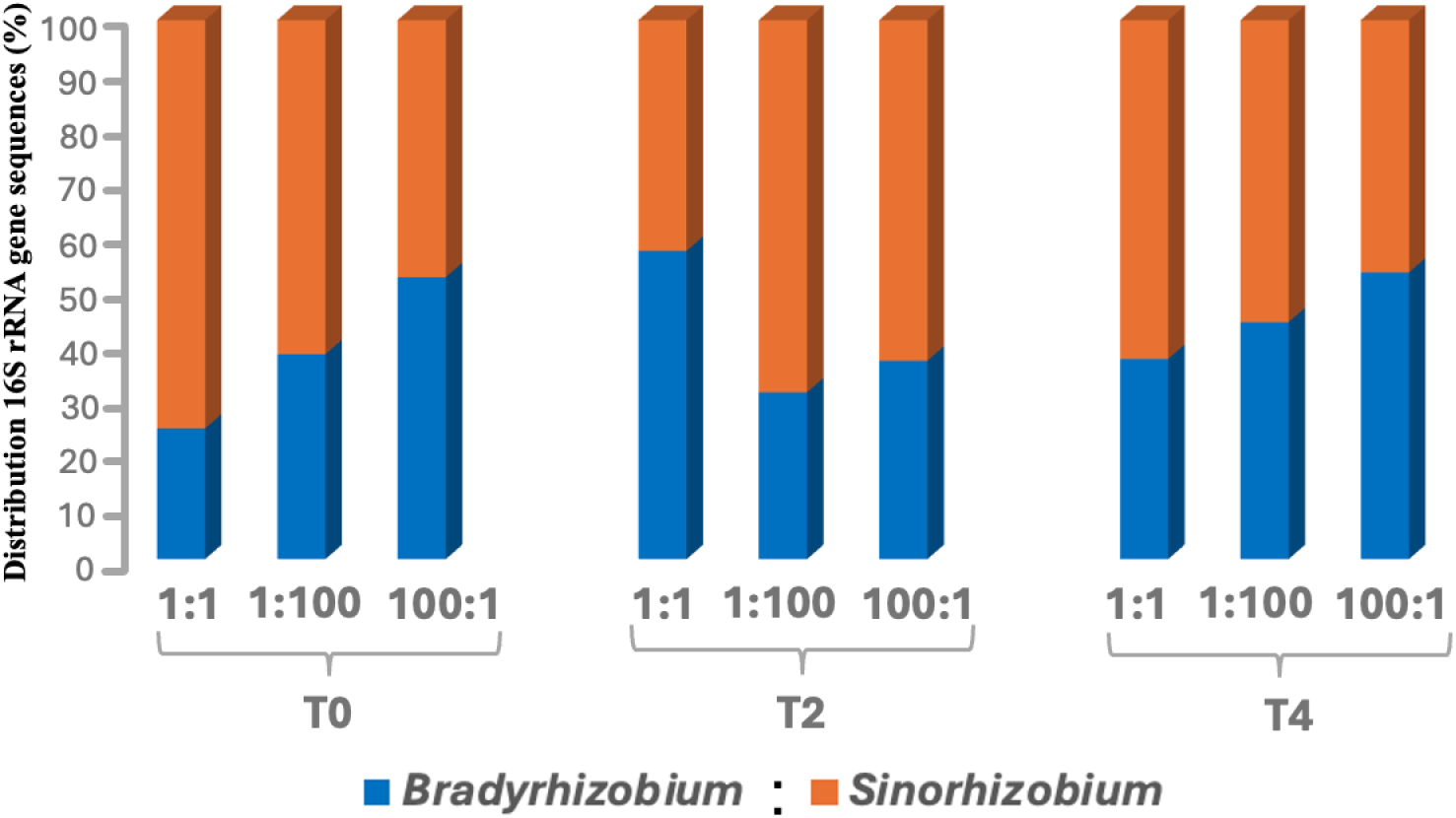
Relative distribution of *Bradyrhizobium diazoefficiens* USDA 110 (BR) and *Sinorhizobium fredii* USDA 191 (SR) sequences in the rhizosphere of soybean plants at harvest (Day 56). Rhizosphere samples were collected from 71 plants co-inoculated with BR and SR at sowing (T_0_), two weeks (T_2_), and four weeks (T_4_) after germination, using inoculation ratios of 1:1, 1:100, and 100:1. A total of 68,173 sequences were retrieved, including 29,351 BR and 38,822 SR sequences.

Although we used sterilized sand at time of seed sowing, sequences affiliated with non-inoculated bacterial genera were consistently detected in the rhizosphere of all plants at the time of final harvesting (8 weeks of plant growth). This indicates cross-contamination through aerial transmission from external sources such as neighboring plants or greenhouse soil. Across all treatments, *Bacillus* was the dominant genus (averaging ∼2,500-18,000 sequences per treatment), followed by *Comamonas* and *Alicyclobacillus*, which occurred at low abundances. A minor fraction of sequences related to other genera such as *Geobacter, Mycobacterium, Symbiobacterium*, and *Acidaminococcus* were also detected (**Fig. 4; Table S1**). *Bacillus* spp. among these non-rhizobial genera have been previously reported to promote the growth of *Sinorhizobium* spp., while suppressing the growth of *Bradyrhizobium* species and ultimately influencing their relative colonization in nodules (16). These non-inoculated genera were consistently detected across all treatments, despite significant differences in BR and SR occupancy within soybean nodules, and thus may not exert as strong an influence on nodule colonization as previously suggested (16).

**Fig. 4.**
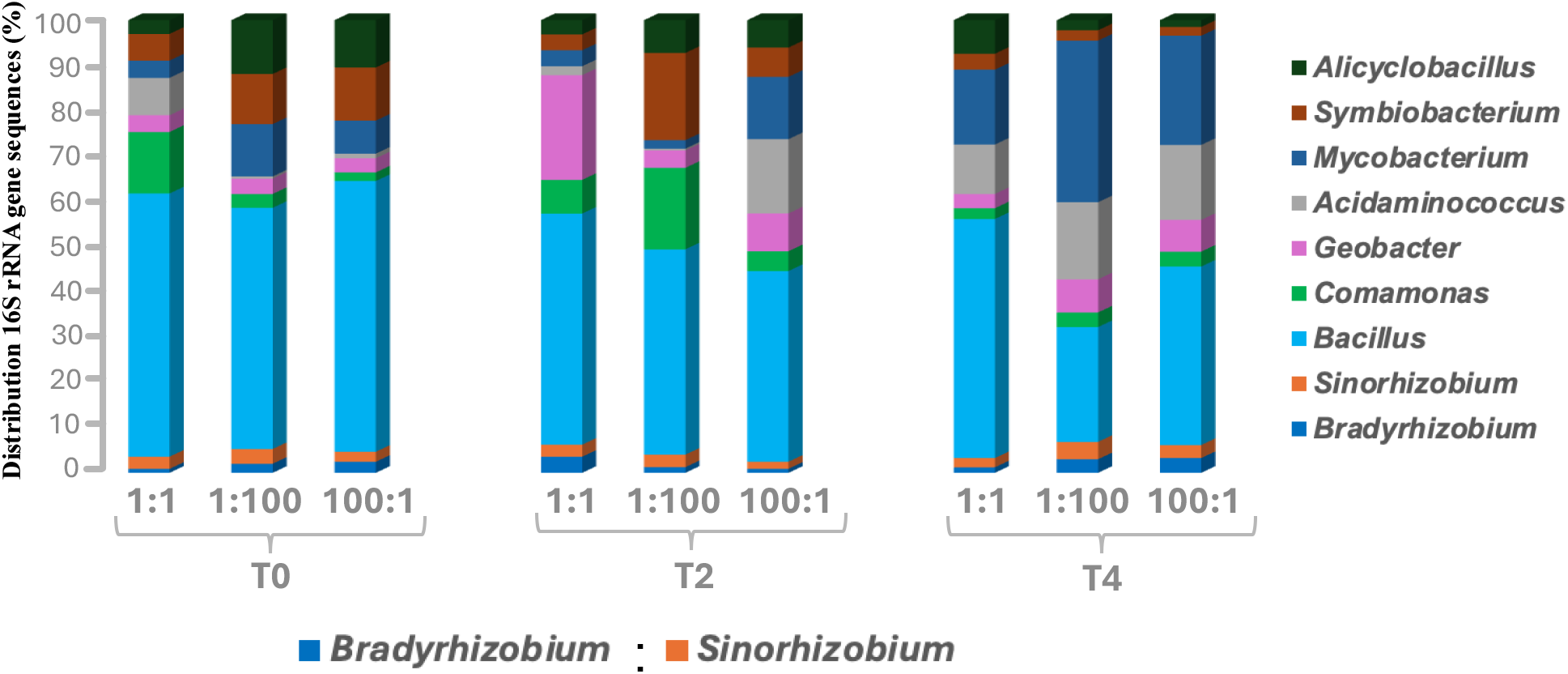
Distribution of bacterial sequences related to rhizobial and non-rhizobial genera in the rhizosphere of 71 soybean plants co-inoculated with *Bradyrhizobium diazoefficiens* USDA 110 (BR) and *Sinorhizobium fredii* USDA 191 (SR). Plants were inoculated at sowing (T_0_), two weeks (T_2_), and four weeks (T_4_) after germination, using BR:SR ratios of 1:1, 1:100, and 100:1. A total of 2.13 million sequences were retrieved from rhizosphere samples.

Although not the primary focus of the current study, we assessed various plant growth characteristics alongside root nodule occupancy by BR and SR strains. Our results indicated no significant trend in plant growth, regardless of whether BR or SR dominated root nodules (**Table 1**). The number of nodules per plant and root–shoot mass per plant remained relatively consistent across treatments. Overall, some fluctuation was observed in specific traits such as root-shoot mass and nodule number; however, these differences were minimal and showed no consistent trend across time points or treatments (**Table 1**).

### Summary

Our study demonstrates critical role for host plant age in the preferential selection between *Bradyrhizobium diazoefficiens* USDA 110 (BR) and *Sinorhizobium fredii* USDA 191 (SR). When inoculated at seed sowing, BR dominated within soybean root nodules. However, as plants matured, SR outcompeted BR, ultimately dominating nodule occupancy at later inoculation stages regardless of inoculum ratio. This age-dependent selection may arise from changes in root exudate composition, signaling molecule profiles, and nitrogen demands of the plant. Despite distinct patterns of nodule occupancy, BR and SR sequences maintained similar relative abundances in the rhizosphere at harvest, indicating that host-driven selection rather than rhizobial availability determines nodule colonization. These findings have significant implications for improving legume inoculation practices, especially under field conditions where the timing of inoculation can be optimized to favor beneficial microsymbiont partner.

## MATERIALS AND METHODS

### Bacteria Culturing and Seed Inoculation

Bacterial cultures of *Bradyrhizobium diazoefficiens* USDA 110 and *Sinorhizobium fredii* USDA 191 were obtained from the ARS Culture Collection (Northern Regional Research Laboratory, Peoria, Illinois). Cultures were grown in yeast mannitol broth (YMB) to achieve comparable cell densities. *B. diazoefficiens* is a slow-growing microorganism and was cultured for 7 days to reach a concentration of 2.38 × 10^7^ CFU/mL. In contrast, *S. fredii* grew more rapidly, requiring only 2 days to reach a similar density of 2.71 × 10^7^ CFU/mL. Colony-forming units (CFUs) were determined using standard plate count methods by serially diluting the cultures and plating them on yeast mannitol agar (YMA) amended with Congo Red dye. For inoculations, both cultures were grown in 250 mL of YMB, then harvested by centrifugation at 10,000 rpm for 5 minutes. The resulting cell pellets were resuspended in 1% phosphate-buffered saline (PBS) and serially diluted in 50 mL tubes. Optical density (OD) was measured using single-beam visible spectrophotometer (Bausch and Lomb Spectronic 20). Cells were diluted to comparable OD values: 0.564 for *B. diazoefficiens* and 0.576 for *S. fredii*, measured at a wavelength of 600 nm. These OD measurements were calibrated with the standard plate count using standard curve generated using triplicate OD readings and CFU counts. A comparable number of cells were harvested for both cultures (2.40 × 10^7^ ± 0.35 CFU/mL). The same procedure was adopted for the later stages of plant inoculation (T_2_ and T_4_) where cells were harvested to similar OD for inoculation.

Different cell ratios of *B. diazoefficiens* and *S. fredii* were prepared by combining defined volumes of cell suspensions in PBS. For a 1:1 ratio, 2.5 mL of each suspension was mixed in a 15 mL conical tube and briefly vortexed. To create a 1:100 ratio, 50 µL of *B. diazoefficiens* was combined with 5 mL of *S. fredii*. For the 100:1 ratio, the same volumes were used in reverse. Single-strain controls were also included: 1:0 for *B. diazoefficiens* only, and 0:1 for *S. fredii* only.

These mixtures were used to inoculate soybean plants at three time points: seed sowing (T_0_), two weeks (T_2_), and four weeks (T_4_) after plant growth. For seed inoculation, soybean seeds were placed into 1-inch deep holes, and 100 µL of inoculum (∼2.40 × 10^6^ ± 0.06 CFU/100 µL) was pipetted directly onto each seed. For T_2_ and T_4_ treatments, 100 µL of the respective inoculum was applied to the base of each plant, followed by 5 mL of sterile water to ensure even distribution of rhizobial cells in the rhizosphere. Likewise, the control plants were inoculated with 100 µL of the respective strain (either BR or SR).

### Greenhouse Study

Soybean seeds of the Agate cultivar were obtained from the Seed Savers Exchange Organization (Decorah, Iowa, USA). Seeds were grown in autoclaved landscaping sand (Sakrete, 0.5 cubic feet) placed in 3 Gallon plastic pots that had been washed with 10% bleach, followed by washing with sterilized distilled water. Prior to sowing, seeds were surface sterilized by sequentially rinsing with 70% ethanol, followed by hydrogen peroxide, and finally rinsed three times with sterile distilled water.

Four holes, approximately one inch deep, were made in each pot, and one sterilized seed was placed in each hole. Soybean plants were grown in a greenhouse under controlled conditions: 28°C temperature, 50–70% relative humidity, and a 14-hour light/10-hour dark photoperiod. During the first week, daily watering was done by surface application (50 mL); however, after seed germination, watering was carried out via subirrigation by placing saucers beneath the pots, allowing water to rise through capillary action (49). Plants were supplemented weekly with half-strength Hoagland solution through surface application. Inoculated plants received a nitrogen-free, half-strength Hoagland solution.

Inoculations were performed at different time points based on plant growth stage. T_0_ plants were inoculated at the time of sowing (September 20, 2024). T_2_ plants were inoculated at two weeks of age (October 4, 2024), and T_4_ plants were inoculated at four weeks after planting (October 18, 2024). A total of 50 pots were used in the experiment. At each inoculation time point, 12 pots were assigned to treatment groups, including four replicate pots for each of the three cell ratios (1:1, 1:100, and 100:1). In addition, two replicate pots were included for each single-strain control, inoculated with either *Bradyrhizobium* or *Sinorhizobium*, and two unplanted pots containing sterile sand served as controls. This brought the total number of pots for the T_0_ (at-sowing) treatment to 18. For the T_2_ (2 weeks post-germination) and T_4_ (4 weeks post-germination) treatments, 16 pots were used at each time point, as the unplanted controls were omitted. Soybeans were grown for 56 days and harvested at the flowering stage. From each pot, four soybean plants were analyzed for root nodule and rhizosphere microbiomes. In total, the microbiome of root nodules from 95 different plants (10 randomly selected nodules per plant) and the rhizosphere microbiome of 71 co-inoculated plants were analyzed. Additionally, we repeated the experiment with a 1:1 inoculation ratio at both T_0_ and T_4_ stages and observed a similar preferential selection (>85% sequences) for SR at the later growth stages (data not shown).

### Harvesting and Collection of Plant Samples

We assessed a few plant growth parameters, such as number of nodules, and root and shoot dry weight as described previously (50). Soybean roots and shoots were separated by cutting the plants with sterile scissors approximately 1 cm above the root crown. Each part was placed into appropriately labeled paper bags. Roots were carefully removed by inverting the pots, and bulk sand was gently dislodged onto sterilized plastic sheets. Rhizosphere sand closely adhering to the roots was collected by gentle shaking and transferred into sterile Eppendorf tubes for downstream bacterial community analysis. Root nodules from each plant were carefully detached using sterile tweezers and placed into separate Eppendorf tubes for endophytic bacterial analysis. In addition to root nodule and rhizosphere sand samples were also collected for bacterial analysis.

### Root Nodule Processing and DNA Extraction

Root nodules were thoroughly washed with sterile distilled water and surface-sterilized using 75% ethanol. The epidermis of each root nodule was aseptically removed using sterilized forceps while the nodules remained submerged in 75% ethanol, following the protocol described previously (14). A total of 10 randomly selected surface-sterilized root nodules per plant were crushed in 1 mL of sterile water using a sterilized mortar and pestle. The resulting homogenate was used for DNA extraction with the DNeasy PowerLyzer PowerSoil DNA Extraction Kit (Qiagen).

### PCR and Sequencing Preparation

To analyze bacterial communities, the 16S rRNA gene was amplified and sequenced from both root nodules and rhizosphere sand. A two-step PCR approach was used as described previously (14). In the first PCR, the V3-V5 region of the 16S rRNA gene was amplified using universal bacterial primers 515-F (5’-GTGCCAGCMGCCGCGG-3’) and 907-R (5’-CCGTCAATTCMTTTRAGTTT-3’). Agarose gel electrophoresis was used to visualize amplicons from each round of PCR. Conditions for the first round of PCR were as follows: Initial denaturation 94°C for 7 minutes, 35 cycles of denaturation at 94°C for 30 seconds, annealing at 56°C for 30 seconds, extension at 72°C for 30 seconds, and a final extension at 72°C for 7 minutes (14). Negative controls ran on gels along with each set of PCR reactions. The first round of PCR samples were cleaned using the ExoSAP-IT PCR cleanup kit (Invitrogen, USA) according to manufacturer’s instructions.

Cleaned products were then used as a template for the second round of PCR. The conditions for the second round were the same except the primers were changed. Primers with two Illumina sequencing adapters (A, 5’-CAAGCAGAAGACGGCATACGAGAT-3’; B, 5’-AATGATACGGCGACCACCGAGATCTACA-3’) were used to target and label samples individually. The conditions for the second round of PCR were as follows: initial denaturation at 94°C for 3 minutes, 15 cycles of 94°C for 30 seconds, annealing at 60°C for 30 seconds, extension at 72°C for 30 seconds, and a final extension at 72°C for 7 minutes (14). PCR products were pooled together, and a final cleanup was performed using an Agencourt AMPure magnetic bead system (Beckman Coulter, Brea, CA). Illumina paired-end DNA sequencing was performed using the NextSeq 2000 platform at the Center for Integrated Biosystems (CIB), Utah State University, USA.

### Data Analysis

Illumina sequencing reads were demultiplexed, and forward and reverse reads were merged. The resulting sequences were denoised and clustered into Amplicon Sequence Variants (ASVs) at 99% similarity using the Quantitative Insights into Microbial Ecology 2 (QIIME 2) pipeline (51). High-quality reads achieved a median Phred score of 30. Taxonomic classification was performed using the SILVA 138.1 reference database, which targets the V4 region of the bacterial 16S rRNA gene. Mitochondrial and chloroplast sequences were subsequently filtered out and removed using QIIME 2.

For statistical analysis, we log-transformed the data. We used the non-parametric Kruskal–Wallis test to compare different cell ratios (1:1, 1:100, 100:1; χ^2^ = 2.82, df = 2, P = 0.2436) and inoculation times (T_0_, T_2_, T_4_; χ^2^ = 19.17, df = 2, P < 0.001). The results indicated that inoculation time had a significant effect on whether *Bradyrhizobium diazoefficiens* (BR) or *Sinorhizobium fredii* (SR) dominated in root nodules. We reported the relative distribution of BR and SR sequences as percentages in the main text (Table 1) and also showed pairwise comparisons of log-transformed data, along with the standard error bars in the supplemental material (**Fig. S3 & S5**). Additionally, we also re-assessed all data after normalizing for 16S rRNA gene copy number for SR (three copies per genome) and reported the results in supplemental material. Lastly, it is important to mention here that nodule sequence data were heavily dominated by one strain or the other, and the distribution of 16S rRNA gene counts were not normally distributed, suggesting the basic assumptions of ANOVA (normality and homogeneity of variance) were violated. We also attempted to use Poisson generalized linear models (GLMs). However, these models also failed to meet the equidispersion assumption, as the data showed extreme overdispersion (>18,000) and zero inflation, which could inflate Type I error. Lastly, using ratios (BR/SR or SR/BR) did not resolve these issues, as many values were either very large or close to zero. All statistical analyses were conducted using R (R Core Team, 2023) (53).

### Nucleotide sequence accession numbers

The 16S rRNA gene sequences were deposited in the NCBI Sequence Read Archive (SRA) under Bio-project PRJNA1338725.

